# Ionizing Radiation induces cells with past caspase activity that contribute to the adult organ in *Drosophila* and show reduced Loss of Heterozygosity

**DOI:** 10.1101/2022.11.18.517019

**Authors:** Sarah Colon Plaza, Tin Tin Su

## Abstract

There is increasing recognition that cells may activate apoptotic caspases but not die, instead displaying various physiologically relevant consequences. We know very little, however, of the mechanisms that underlie the life/death decision in a cell that has activated apoptotic caspases. By optimizing a published reporter for past caspase activity, we were able to visualize cells that survived caspase activation specifically after exposure to ionizing radiation in *Drosophila* larval wing discs. We found that cells with <u>X</u>-ray-induced <u>p</u>ast <u>a</u>ctive <u>c</u>aspases (XPAC) do not arise at random but are born at specific locations within the developing wing imaginal discs of *Drosophila* larvae. We found also that the apoptotic signaling pathway is needed to induce XPAC cells. Yet, XPAC cells appear in stereotypical patterns that do not follow the pattern of IR-induced apoptosis, suggesting additional controls at play. Functional testing identified the contribution of *wingless* (*Drosophila* Wnt1) and Ras signaling to the prevalence of XPAC cells. By following irradiated larvae into adulthood, we found that XPAC cells contribute to the adult wing. To address the relationship between XPAC and genome stability, we combined a reporter for past caspase activity with *mwh*, an adult marker for Loss of Heterozygosity (LOH). We found a lower incidence of LOH among XPAC compared to cells that did not activate the reporter for past caspase activity. In addition, at time points when wing disc cells are finishing DNA repair, XPAC cells show an anti-correlation with cells with unrepaired IR-induced double-stranded breaks. Our data suggest that non-lethal caspase activity safeguards the genome by facilitating DNA repair and reducing LOH after transient exposure to X-rays. These results identify a physiological role for non-lethal caspase activity during recovery from radiation damage.

## Introduction

An exciting emerging field is based on the recognition that cells may activate apoptotic caspases but not die, instead displaying various physiologically relevant consequence such as axon pruning and reduction of the cytoplasm during sperm maturation (for reviews, (1-4)). In one variation, a cell may return to life from the brink of apoptotic death in a process called anastasis (2, 5-7), which is seen in cultured cells exposed to a variety of death stimuli including ethanol (8), a broad-spectrum kinase inhibitor Staurosporine and a small molecule BCL-2 antagonist (9), or ion beam radiation (10). Removal of death-inducing stimuli before apoptosis is completed allows some cells to recover even after displaying such apoptotic features as cleaved caspases, Mitochondrial Outer Membrane Permeabilization, or phosphatidylserine flipping. Cells that underwent anastasis are not only alive but are capable of proliferation and initiating tumors. How cells activate apoptotic caspases but not die is still poorly understood. Dissecting this phenomenon requires the ability to detect cells that activated caspases but did not die. Two genetic reporters for past caspase activity have been described in *Drosophila melanogaster* (11, 12). CasExpress and Caspase Tracker relies on the transcriptional activator GAL4 with a cell-membrane tether that includes a recognition sequence, DQVD, for effector caspase Drice (*Drosophila* caspase 3/7 in conjunction with the G-trace lineage tracing system (13)). Upon Drice activation, the tether is cleaved and GAL4 enters the nucleus to initiate the expression UAS-FLP recombinase, which catalyzes a recombination event to cause permanent GFP expression. Both sensors show widespread non-lethal caspase activity in nearly every tissue during *Drosophila* development (11, 12). CasExpress reporter was used in a screen for regulators of developmental non-lethal caspase activity that identified apical caspase Dronc, caspase inhibitor p35, pro-survival signaling component Akt, transcription factor dCIZ1 and Ras (14). One of these, dCIZ1, was shown to promote survival after caspase activation by heat shock, oncogenic stress and X-rays.

We wanted study cells with past caspase activity induced not by developmental events but by ionizing radiation (IR). IR is used to treat more than half of cancer patients where therapeutic success relies on IR’s ability to induce apoptosis. Therapy failure can result if cells initiate but do not complete apoptosis after IR exposure, making it important to understand molecular mechanisms that regulate the production of such cells. The use of CasExpress and Caspase Tracker to detect IR-induced non-lethal caspase activation have been challenging because of strong developmental signals from these biosensors even when GAL4 activity is temporally controlled with the temperature sensitive repressor GAL80^ts^ (15). Here, we report an optimized temperature shift protocol that out-performs published protocols by increasing the ratio of IR-induced to constitutive signal from ∼2-4 to ∼20. This allows us to detect cells with past caspase activity that results specifically from exposure to X-ray, a type of IR. We found that cells with <u>X</u>-ray-induced <u>p</u>ast <u>a</u>ctive <u>c</u>aspases (XPAC) do not arise at random but are born at specific locations within the developing wing imaginal discs of *Drosophila* larvae. Inhibition of apoptotic signaling reduced the prevalence of XPAC cells. Yet, XPAC cells appear in stereotypical patterns that do not follow the pattern of IR-induced apoptosis, suggesting additional controls at play. Functional testing identified Wg (Wnt1) pathway and Ras signaling as such controls. By following irradiated larvae into adulthood, we found that XPAC cells contribute to the adult wing. Anastasis has been associated with increased genome instability in immortalized mammalian cells (7, 9, 10). On the other hand, wide-spread non-lethal caspase activity is detected during normal *Drosophila* development, raising the possibility that such activity may be neutral or even beneficial to healthy cells. XPAC cells, we found, show reduced Loss of Heterozygosity (LOH), suggesting that non-lethal caspase activity safeguards the genome after transient exposure to X-rays in *Drosophila*.

## Results

### Optimizing temperature shift conditions to detect cells with past ionizing radiation-induced caspase activity

In our published work with Caspase Tracker, we raised the larvae at 18°C and shifted them to 29°C to inactivate GAL80^ts^ from 0 to 6h after exposure to 4000R (40Gy) of X-rays (15) (Fig. 1A, T=29°C). 4000R is typically used in *Drosophila* because it kills more than half of the cells in a wing disc but is still compatible with regeneration and survival as viable fertile adults can emerge. We chose a 6h temperature shift based on the literature. *hid* and *rpr*, which encode SMAC orthologs and are essential for IR-induced caspase activation, are transcriptionally induced at 2h after IR (16). Caspase cleavage and staining with the vital dye Acridine Orange (AO) are detectable at 3h after IR; AO stains apoptotic and not necrotic cells in *Drosophila* (17). By 4h after IR, *hid/rpr* expression has declined compared to the 2h time point, although cleaved caspases and AO staining remain visible for up to 24h after (16). Based on these results, a 6h window immediately after IR would capture the bulk of IR-induced caspase activation.

**Figure 1.**
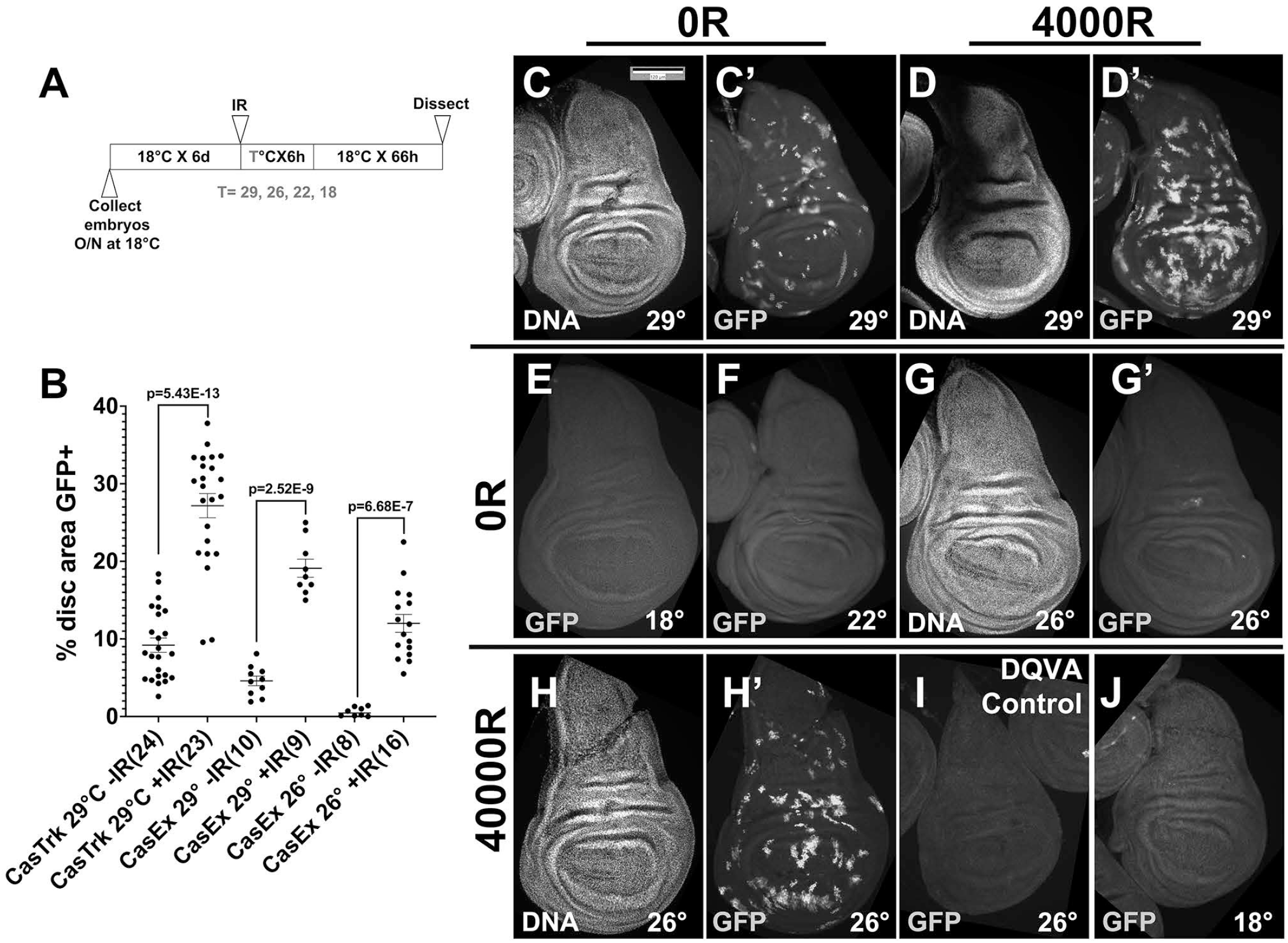
Optimizing temperature shift conditions to detect cells with X-ray-induced past caspase activity. Larvae were treated as in Fig. 1A where T= 29°, 26°, 22° or 18°C as indicated on the panels. Wing discs were dissected at 72h after exposure to 0R (-IR) or 4000R (+IR), fixed, stained for DNA, and imaged for DNA and GFP. CasTrk = Caspase Tracker. CasEx = CasExpress. The genotype of the wing discs shown was CasExpress/G-trace; GAL80^ts^/+ in all panels except (I) which shows a control CasExpress mutant with a DQVD>DQVA mutation that renders the GAL4 membrane tether caspase-resistant. Scale bar = 120 microns. (A) The experimental protocol (B) Fractional area of each disc that is GFP+ was quantified in Image J. p-values were calculated using a 2-tailed t-test. The numbers in the brackets are the number of discs analyzed in two to four biological replicate experiments per sample. (C-C’) An unirradiated disc shows GFP+ cells even after a short (6h) pulse of heat inactivation of GAL80^ts^ at 29°C. (D-D’) An irradiated disc shows more GFP+ cells with the same temperature shift as in C. (E-G’) Unirradiated discs show few GFP+ cells when the temperature shift for 6h is to 18°C (E), 22°C (F) or 26°C (G-G’). (H-H’) A irradiated discs from larvae shifted to 26°C shows robust GFP signal. (I) Discs from caspase-resistant CasExpress mutant larvae treated exactly as in H. (J) Discs from larvae treated exactly as in (H) but with no temperature shift

Even with limited (6h) activation of GAL4, unirradiated wing discs from unirradiated larvae show GFP+ cells (Fig. 1C’, quantified in Fig. 1B, 29° -IR datasets), as we reported before [20]. Although the GFP+ area increased significantly with irradiation by 3-fold for Caspase Tracker and 4-fold for CasExpress (Fig. 1D’, quantified in B), high constitutive signal means about 66% of Caspase Tracker>GFP cells and about 75% of CasExpress>GFP cells in irradiated discs are X-ray-induced while the remainder are from developmental caspase activation. To eliminate the GFP signal from the latter, we tried using lower temperatures to inactivate GAL80 in the protocol in Fig. 1A (Figure 1 E-G’), using CasExpress that shows smaller variance than Caspase Tracker -IR (Fig. 1B). We found that we could go as high as 26°C and still keep the CasExpress>GFP area to near zero in unirradiated discs (Fig. 1G’, quantified in B) while IR increased the GFP+ area significantly (Fig. 1H’, quantified in B). A control CasExpress sensor with a DQVD->DQVA mutation showed no GFP+ cells even with IR (compare Fig. 1I to H’), indicating that GFP+ cells report effector caspase activity specifically. Returning the larvae back to 18°C immediately after irradiation, that is, without any temperature shift, eliminated GFP+ cells (Fig. 1J), indicating that production of GFP+ cells is GAL4-dependent. Because of low GFP+ area without IR at 26°C, the ‘signal to noise’ ratio increased to 20-fold, meaning most (>90%) of GFP+ cells +IR discs were cells with X-ray-induced past active caspases (XPAC). Subsequent experiments use the protocol in Fig. 1A with T=26°C.

### XPAC *cells proliferate*

The data in Fig. 1 were from 3 days after irradiation. To study GFP+ XPAC cells soon after they form, we performed a time course (Fig. 2). GFP+ cells were not observed at 6h after IR but were readily detectable at 24h after IR (Fig. 2A). At this timepoint, most GFP+ cells were either singlets or in small groups of 2-5 cells. These results agree with the published timings of check point arrest, recovery and cell doubling in the wing disc. 4000R of X-rays stops the cell cycle via checkpoint activation, with S phase and mitosis resuming at about 6h after irradiation (18). Cell doubling times in 3^rd^ instar wing discs average 12h (19). Therefore, at 24h after irradiation, cells would have divided on average just once, which can be seen as the appearance of singlets (arrow) and doublets (arrowhead) in Fig. 2A. At 48h after irradiation, single GPF+ cells were rare and most cells were in clones of 4-30 cells each (Fig. 2B). Clone size increased further at 72h after IR, the last time point taken before we lost the larvae to pupariation (Fig. 2C). We conclude that XPAC cells are capable of clonogenic proliferation. Real time marker RFP was not observed presumably because the return to 18°C restored GAL80^ts^ and repressed GAL4.

**Figure 2.**
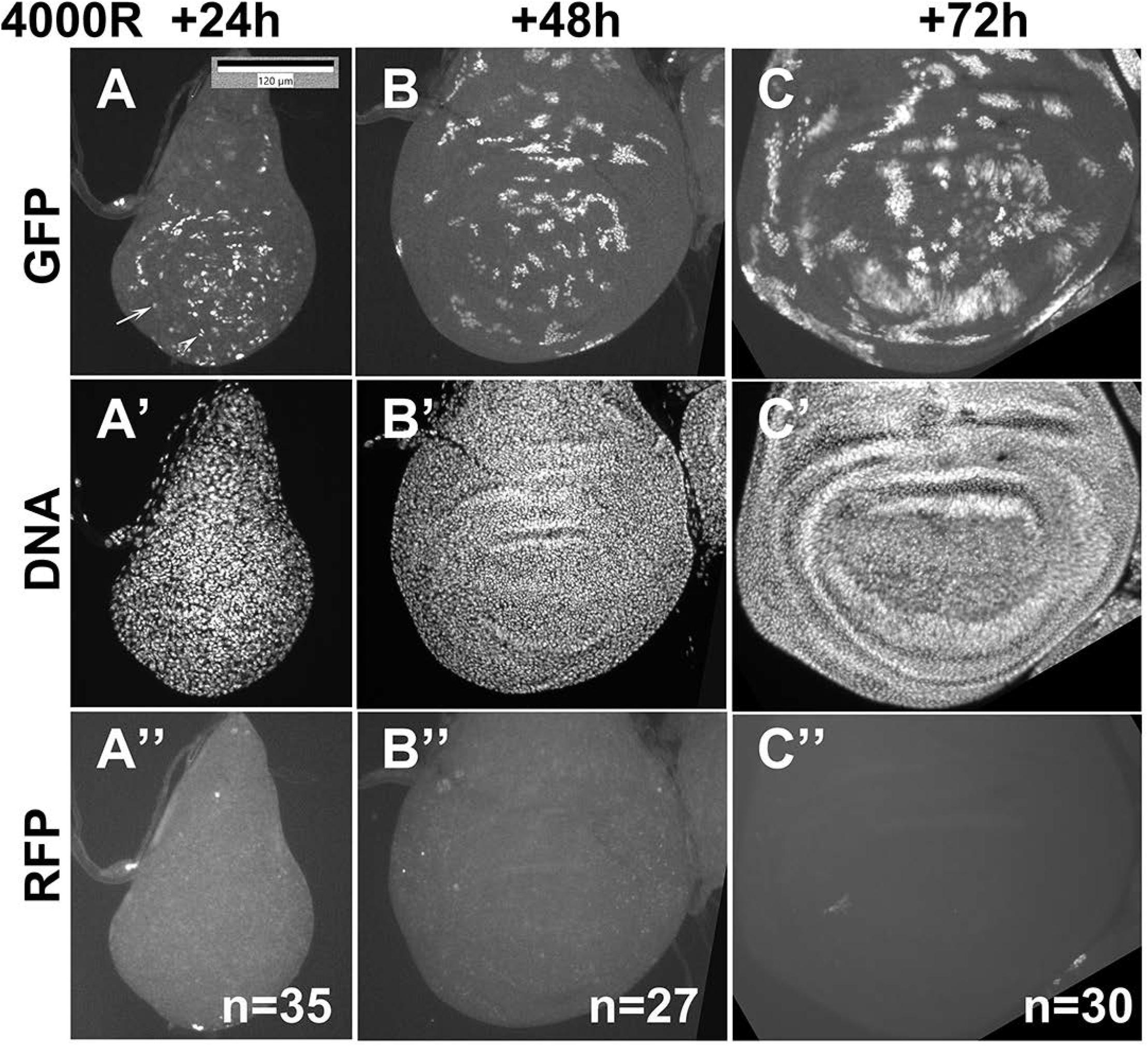
XPAC cells are capable of proliferating. Larvae of the genotype CasExpress/G-trace; GAL80^ts^/+ were treated as in Fig. 1A where T=26°C. Wing discs were dissected at 24, 48 and 72h after exposure to 4000R, fixed, and stained for DNA. The discs were imaged for GFP (A-C), DNA (A’-C’) and RFP (A’’-C’’). N= number of discs examined in three biological replicates for 24h and 2 biological replicates each for 48 and 72h. The arrow points to a singlet GFP+ cell and the arrowhead points to a doublet of GFP+ cells in panel A. Scale bar = 120 microns.

### The prevalence and the location of XPAC cells change with development

Our collection of larvae spanned 24h in age and show a range of developmental stages, that is, wing discs of different sizes and degree of epithelial folding (Figure 3A-D, arrowheads). We noticed three trends in the prevalence and the location of GFP+ XPAC cells at 24h after IR. First, smaller discs showed more GFP+ area relative to the disc size than larger, more developed discs (compare Fig. 3D’ to A’). A plot of % disc area, that is GFP+ against disc size illustrates this trend (Fig. 3F ‘6-7d at IR’). The area serves as surrogate for cell number since wing discs are composed of a single layer epithelium. We quantified clone area instead of clone number to avoid errors in assigning cells to clones, especially as XPAC cells appear close together (Fig 2A-C). By using an early (24h) time point, we are catching XPAC cells soon after birth and before cell division makes a significant contribution to clone area (see preceding paragraph). We interpret the dependence of XPAC prevalence on disc size to mean that as the disc increased in developmental age, X-rays were able to induce fewer cells that activated caspases but survived. This correlation was confirmed by aging the larvae for an additional 24h at 18°C before irradiation. This produced larger, more developed discs. Importantly, the same relationship between disc size and % GFP+ area was seen in the older cohort (Fig. 3F, ‘7-8d at IR’), except that overall values shifted to the right as expected for older discs. Separating discs based on the absence or presence of epithelial folds, a criterion used before to separate discs according to development (20), led to the same conclusion; more developed discs had fewer XPAC cells (Fig. 3G).

**Figure 3.**
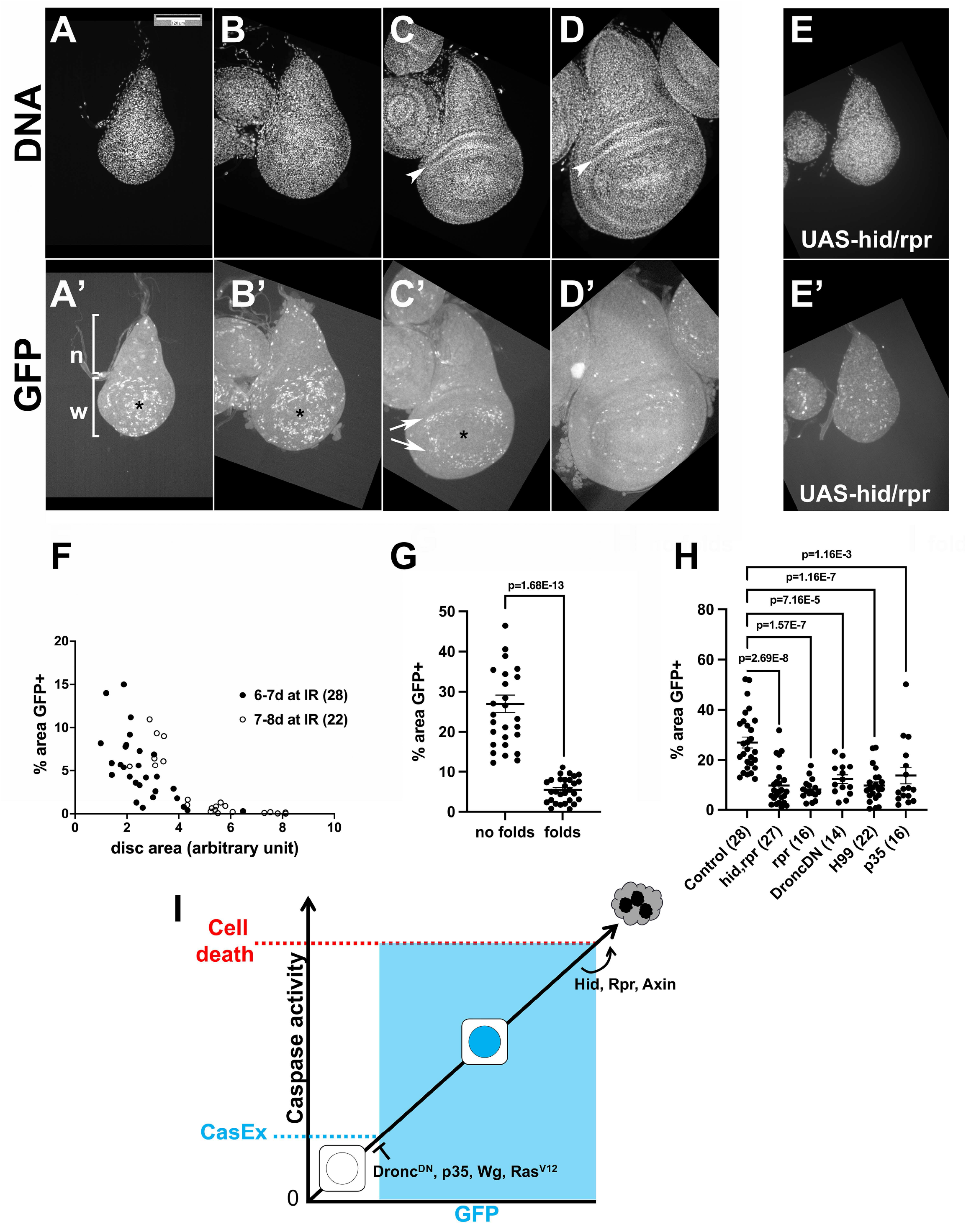
XPAC prevalence in development and the effect of apoptotic signaling. Larvae of the genotype CasExpress/G-trace; GAL80^ts^/+ were treated as in Fig. 1A where T=26°C. Wing discs were dissected at 24h after exposure to 4000R, fixed, and stained for DNA. Scale bar = 120 microns. p-values were calculated using a 2-tailed t-test. (A-D’) The discs were imaged for DNA (A-D) or GFP (A’-D’). Hinge folds are visible in DNA images of more developed discs and are indicated with arrowheads in (C-D). n = notum and w = future wing that includes the hinge and the pouch. The most distal region of the future wing is indicated with * in (A’-C’); GFP+ cells clear from this region before the rest of the future wing. A ring of GFP+ cells is indicated with arrows (C’). (E-E’) CasExpress/G-trace; GAL80^ts^/+ larvae that also carried one copy of UAS-hid, rpr on the X chromosome were treated as in (A-D). (F) The % of disc area that is GFP+ was quantified from discs such as those shown in A’-D’ and plotted against the total disc area. The numbers of discs analyzed are in parentheses. The data are from 3 (6-7d) and 2 (7-8d) biological replicates each. (G) The % of disc area that is GFP+ was quantified for discs with no epithelial folds (less developed discs such as those in A and B) and discs with epithelial folds (more developed discs such as those in C and D). The data are from 6 biological replicate samples of larvae 7-8d old at irradiation. (H-I) The % of disc area that is GFP+ was quantified from discs such as those shown in E’ and plotted against the total disc area. The number of discs analyzed are in parentheses. The genotypes were: Control = CasExpress/G-trace; GAL80^ts^/+ hid, rpr = UAS-hid, UAS-rpr/+ or Y; CasExpress/G-trace; GAL80^ts^/+ rpr = UAS-rpr/+ or Y; CasExpress/G-trace; GAL80^ts^/+ DroncDN = CasExpress/G-trace; GAL80^ts^/UAS-Dronc^DN^ H99 = CasExpress/G-trace; GAL80^ts^/H99 p35 = CasExpress/G-trace; GAL80^ts^/UAS-p35

Second, the distribution of GFP+ cells in each disc was not homogenous. The notum (future body wall or ‘n’ in Fig. 3A’) had fewer GFP+ cells than the future wing (‘w’ in Fig. 3A’ includes the hinge and the pouch). This was true regardless of disc size. Third, when the number of GFP+ cells declined in larger discs, their loss began in the distal pouch (* in Fig. 3A’, B’, C’), leaving a ring of GFP+ cells (Fig. 3C’, arrows).

### Apoptosis signaling can increase or decrease XPAC prevalence

We next investigated the role of apoptotic signaling in XPAC cells. Using UAS-transgenes, we increased apoptotic signaling with *hid* and *rpr* together or *rpr* alone and decreased apoptotic signaling with a dominant negative version of apical/initiator caspase Dronc (apical caspase) or p35 (effector caspase inhibitor). This means only in cells that have initiated caspase activation and translocated GAL4 into the nucleus as part of the CasExpress system would express the transgenes cell-autonomously, identifying what keeps cells alive <u>after</u> they began to activate apoptotic caspases. We used also heterozygotes for the H99 chromosomal deficiency that removes *hid, rpr* and *grim* and has been shown to reduce and delay IR-induced apoptosis (16); the effect of H99 would be throughout the organism and not just cell-autonomous or conditional. We analyzed discs with no folds because their robust XPAC incidence (∼30%) provides room to detect changes up or down. Both, increasing and decreasing apoptotic signaling reduced XPAC prevalence (Fig 3H). The seemingly paradoxical result can be explained by the threshold model wherein active caspases cleave and activate other caspases (21, 22), thus amplifying caspase activity to reach a certain level needed for cell death (23). We propose a second, lower threshold for successful recombination and permanent GFP expression (Fig 3I). Here, initial Drice activity translocates GAL4 into the nucleus but Dronc^DN^ and p35 become expressed to prevent the amplification needed to reach the threshold for successful recombination and permanent GFP expression. H99 would reduce effector caspase activity, preventing some cells from activating CasExpress, thus also reducing the number of GFP+ cells. At the other end of the continuum, increasing effector caspase activity with *hid*/*rpr* would push cells over the threshold for cell death, reducing the number of GFP+ cells that activated apoptotic caspases but lived. In sum, we propose that decreasing or increasing apoptotic signaling will keep a cell out of the intermediate zone of caspase activity needed to produce a GFP+ cell.

### Developmental changes in XPAC induction does not correlate with apoptosis or cell proliferation

We next addressed why the prevalence of GFP+ cells decline in more developed discs. One possibility is cell cycle exit; non-dividing cells are known to be refractory to apoptosis, for example in *Drosophila* eye disc (24). Unlike in the eye disc, however, cells in the wing disc continue to proliferate throughout larval stages except for 4-cell wide Zone of Non-proliferating Cells (ZNC) along the dorsoventral boundary in the late 3^rd^ instar wing disc that arrest in G1/G2 (25). Cell cycle exit is unlikely to explain the reduction in XPAC prevalence with developmental stage for the two reasons. First, at 24h after IR when cells have already recovered from checkpoint arrest (18), antibody staining for phospho-Histone H3, a mitotic marker, showed robust and wide-spread mitoses including in most developed discs in our samples where we see no XPAC cells (Fig 4C-C’’). Specifically, we noted robust mitotic activity in the notum (arrows) and the pouch (between dashed lines) that includes the future ZNC. Second, the notum and the pouch have similar cell doubling times (20), yet, as the wing disc develops, XPAC cells are lost from the notum before they are lost from the pouch (Fig 4B’’). We conclude that cell cycle exit is unlikely to be the cause for XPAC reduction that accompanies development.

**Figure 4.**
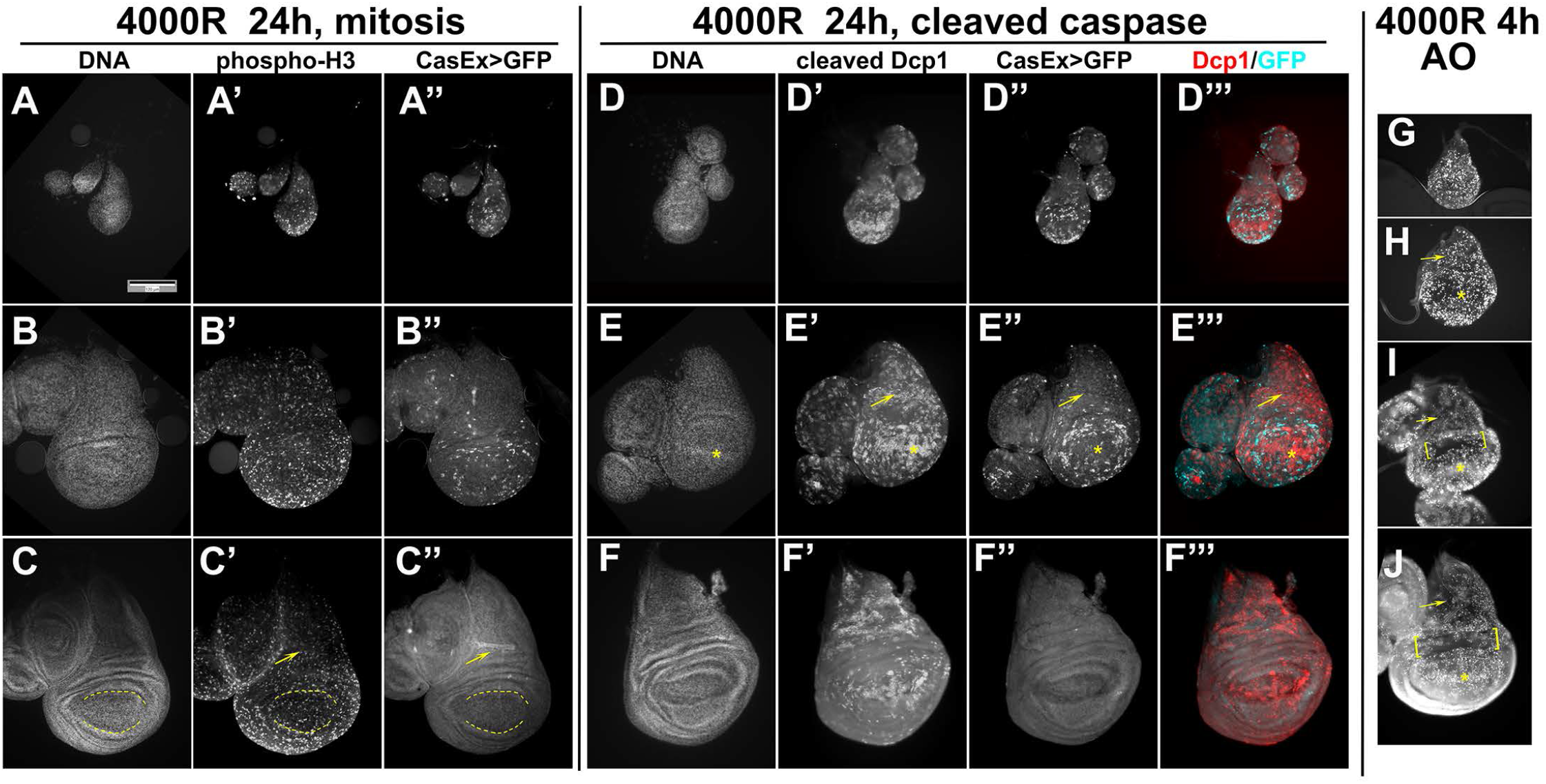
Apoptosis and mitosis are widespread throughout the wing disc. The larvae of the genotype CasExpress/G-trace; GAL80^ts^/+ were treated as in Fig. 1A where T=26°C. Scale bar = 120 microns. (A-C’’) Wing discs were fixed and stained with an antibody to phospho-S10 on Histone H3, a mitotic marker. The dashed line marks the pouch. The discs were also stained for DNA and visualized for CasExpress>GFP. The arrows point to the notum and the dashed lines mark the pouch. (D-F’’’) Wing discs were fixed and stained with an anti-cleaved-Dcp1 antibody to detect cells that activated apoptotic caspases. The discs were also stained for DNA and visualized for CasExpress>GFP. The arrows point to the notum and the * marks the distal pouch. (G-J) Wing discs were stained live with the vital dye Acridine Orange which reports on apoptotic cells. The brackets mark the hinge region that is resistant to IR-induced apoptosis. The arrows point to the notum and the * marks the distal pouch.

The possibility is that X-rays were less efficient at inducing caspase activity in more developed discs is also unlikely. Cleaved active Dcp1 is absent without irradiation (S1 Figure) but is readily discernable in irradiated discs regardless of whether they show robust XPAC induction (Fig 4D’’ and E’’) or only a few detectable GFP+ cells (Fig 4F’’). The notum (arrow) and the distal pouch (*) in particular show robust Dcp1 signal yet display reduced XPAC prevalence as the discs mature. AO staining 4 h after IR confirmed that the induction of wide-spread cell death at developmental stages we study (Fig 4G-H; the brackets indicate the hinge cells that we showed before are refractory to IR-induced apoptosis (24)). We conclude that while apoptotic caspase activity is necessary to generate XPAC cells (Fig 3), it is not sufficient to explain the spatiotemporal distribution of XPAC cells, which does not match the spatiotemporal distribution of IR-induced caspase activation. We infer that other mechanisms operate on top of caspase activation to decide whether a cell with active caspases live or die.

### Wg and Ras Signaling modulate XPAC prevalence

The changing pattern of XPAC cells as discs develop suggests that developmental signaling pathways may play a role. We focused on Wg (Wnt1) and Ras because they are (1) active in the regions of the wing disc with dynamic XPAC prevalence and (2) are known to influence cell survival. For example, XPAC cells clear from the distal pouch around a line of Wg-expressing cells (yellow arrowhead in Fig 5C) while they persist near the Wg Inner Ring (white arrowhead in Fig 5C). As for apoptotic signaling in Fig 3, we used UAS-transgenes to express Wg, Axin (Wg inhibitor) or RAS^G12V^ (constitutively active RAS) specifically in CasEx-GAL4-active cells for 6h after IR. All three transgenes significantly lowered the XPAC prevalence (Fig 5E-F), which is reminiscent of how both increased and decreased apoptotic signaling lowered XPAC prevalence (Fig 3H). Lower XPAC prevalence we see after inhibition of Wg is unlikely to be indirect via delayed development because less developed discs show more XPAC. Given the known anti-apoptotic role of Wg (24), the model in Fig 3I could explain the Wg/Axin results. Specifically, inhibiting Wg with Axin could drive cells over the threshold for cell death while increased Wg could keep the cell below the threshold for CasEx>GFP. Staining for cleaved active Dcp1 supports this idea (Fig 5G); Wg inhibited while Axin promoted caspase activation. The threshold model can also explain the results with RAS^G12V^ (Fig 5F). Ras/MAPK signaling is known to inhibit caspase activation, by direct phosphorylation and inhibition of Hid during *Drosophila* eye development, for example (26). Inhibition of Drice activity by RAS^G12V^ could keep the cell below the threshold for CasEx>GFP, lowering XPAC prevalence. RAS^G12V^ reduced cleaved Dcp1 signal, supporting this idea.

**Figure 5.**
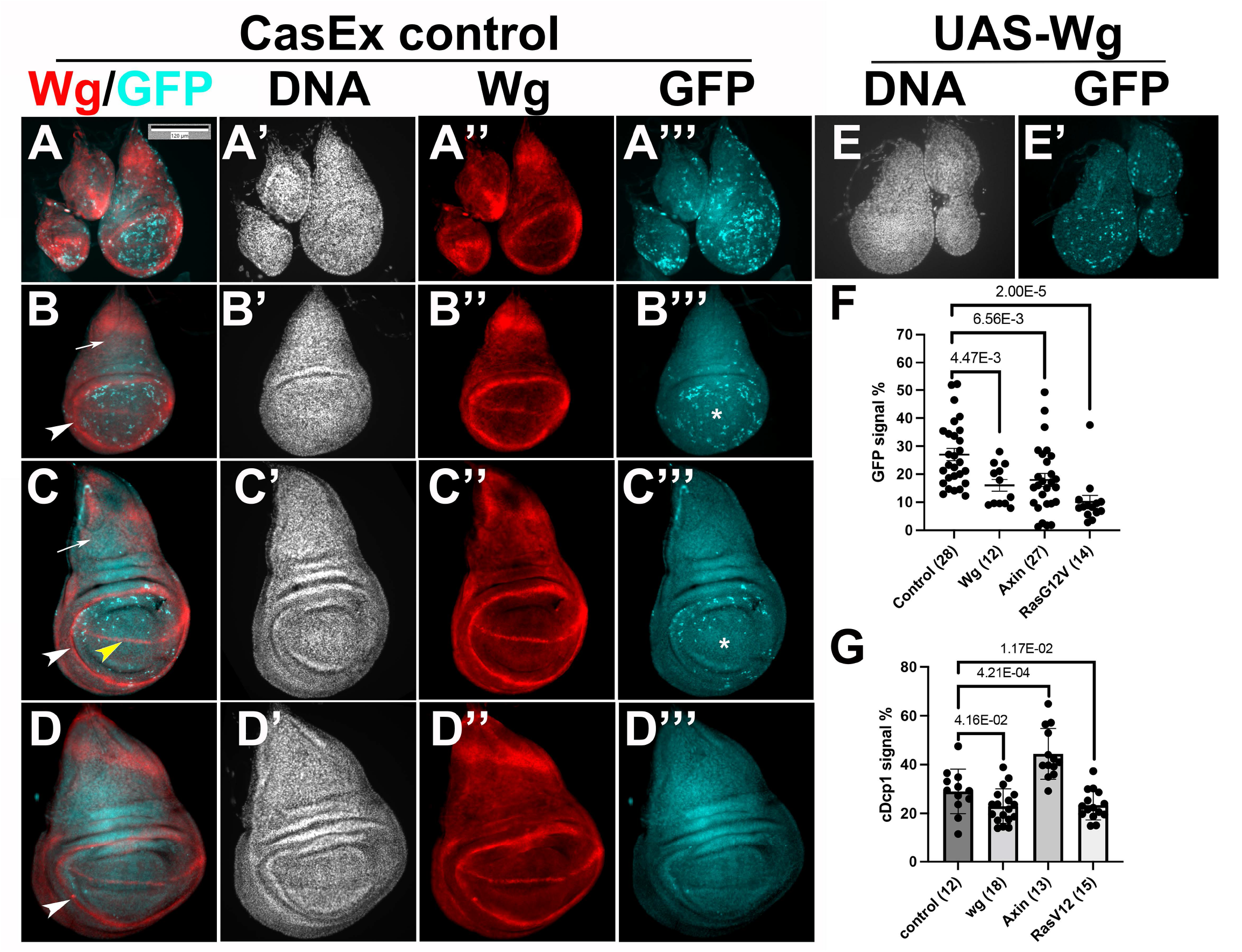
Developmental signaling pathways regulate XPAC cell prevalence. (A-E’) Larvae of the genotypes indicated below were treated as in Fig. 1A where T=26°C. The discs were dissected 24h after exposure to 4000R, fixed and stained with an antibody against Wg (A-D’’’). The discs were also stained for DNA and visualized for CasExpress>GFP. Arrows indicate the notum and * indicate the distal pouch. Arrowheads indicate the Wg Inner Ring (white) or the D/V boundary (yellow). (F-G) The GFP+ areas were quantified and expressed as % of the total disc area. p-values were calculated using a 2-tailed t-test. Scale bar = 120 microns. The genotypes were: Control = CasExpress/G-trace; GAL80^ts^/+ wg = CasExpress/G-trace; GAL80^ts^/UAS-wg Axin = CasExpress/G-trace; GAL80^ts^/UAS-Axin NotchICD = CasExpress/G-trace; GAL80^ts^/UAS-N^ICD^ mam-delta = CasExpress/G-trace; GAL80^ts^/UAS-mam^Δ^ RasV12 = CasExpress/G-trace; GAL80^ts^/UAS-Ras^V12^

### Cells with past caspase activity show reduced IR-induced LOH

We next addressed the consequences of past IR-induced caspase. Cultured human cells exposed to 0.5 Gy (50R) of particle radiation, a type of IR, show persistent caspase activation for up to two weeks without dying, elevated DNA breaks and oncogenic transformation (10). X-rays on the other hand induce transient caspase activation, with DNA breaks clearing within 24h post IR exposure in larval wing discs (18, 24). To assay for long term effects of non-lethal caspase activity under these conditions, we followed XPAC cells into adult wings where they appear in an IR-dependent manner (Fig. 6) regardless of larval age at the time of irradiation (6-7d or 7-8d; corresponding to larval XPAC data in Fig. 3F). This allows us to use the *multiple wing hair* adult marker to quantify Loss of Heterozygosity (LOH), a type of genome aberration. LOH of recessive *mwh* alleles results in homozygous mutant cells with more than one hair each due to defective actin organization (27). We showed previously that *mwh* LOH is absent without IR but is induced when larvae are irradiated with 4000R of X-rays (28). At appropriate magnification, it is possible to assign GFP and *mwh* status to each cell (Fig. 6H). We found that *mwh* incidence was lower for GFP+ relative to GFP-areas of the same wing (Fig. 6I).

**Figure 6.**
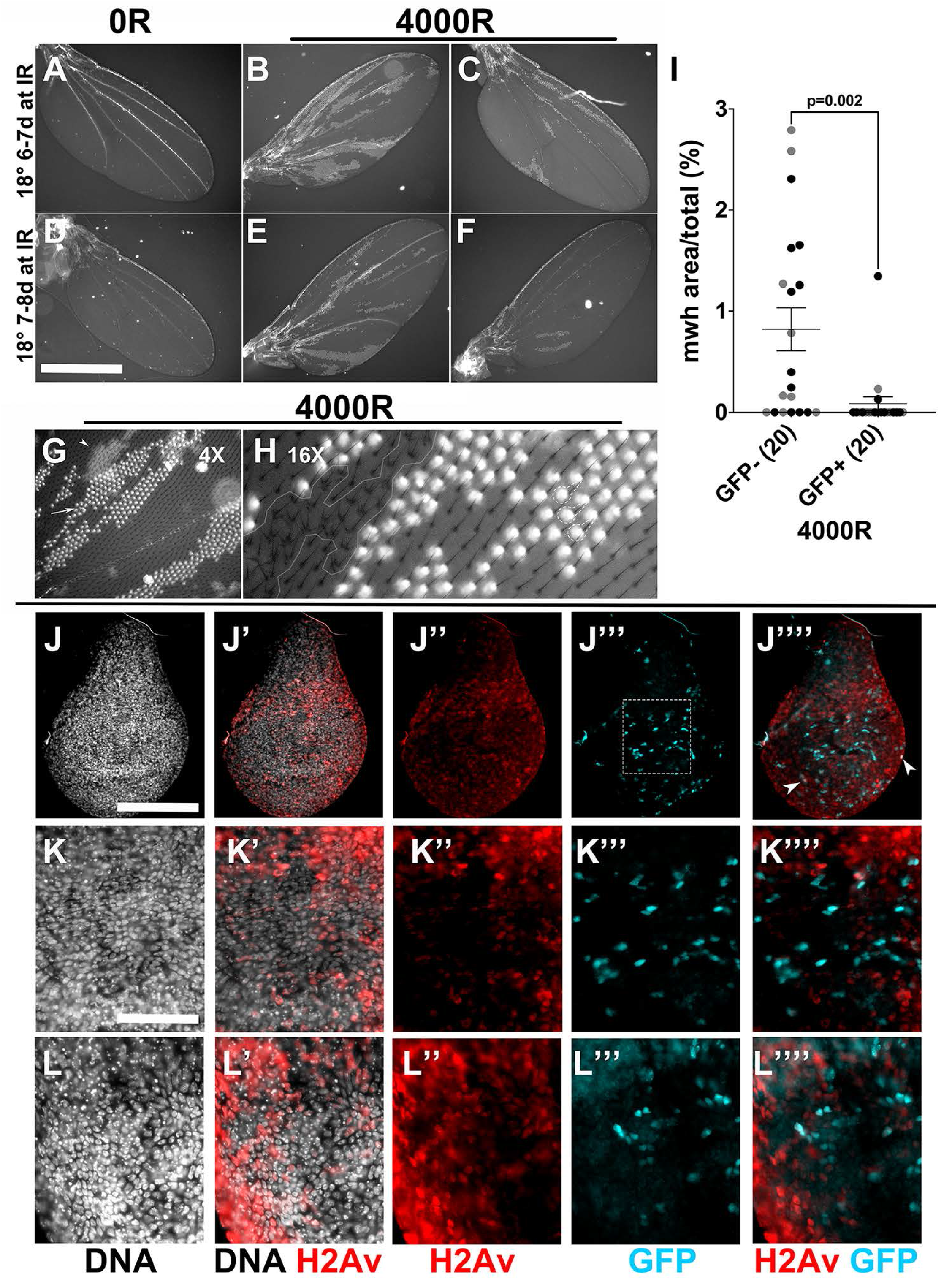
Cells with past caspase activity show reduced IR-induced LOH. (A-F) Adult wings from larvae of the genotype CasExpress/G-trace; GAL80^ts^/mwh^1^. The experimental protocol shown in Fig. 1A was followed but larvae were aged for 6d at 18°C after egg collection to get 6-7d old larvae (A-C) or aged for 7d at 18°C after egg collection to get 7-8d old larvae at the time of irradiation. Larvae were irradiated with 0 or 4000R of X-rays. After incubation at 26°C, larvae were returned to 18°C for the remainder of the experiment. The two +IR wings shown for each group represent the range of clone size and number observed in two biological replicate experiments. The number of wings examined was 12 (6-7d -IR), 9 (6-7d +IR). 6 (7-8d -IR) and 11 (7-8d +IR), in two biological replicate experiments each. (G) A region of the wing was imaged at 4X magnification relative to A-F. It is possible to discern the top (in focus, arrow) and the bottom (out-of-focus, arrowhead) surfaces of the wing in such images. (H) A region of the wing is shown magnified 16X relative to A-F. It is possible to assign mwh state to individual GFP+ cells, for example, 3 cells within dashed lines. In a patch of mwh cells (white line), none are GFP+. (I) Percent of image area that is mwh was quantified from images such as G, for the GFP+ and GFP-portion of the surface in focus. The data include wings from animals irradiated as 6-7d (gray) or 7-8d (black) old larvae. Each dataset also shows a significant difference in mwh between GFP- and GFP+ areas: p= 0.04 for 6-7d and 0.03 for 7-8d. p-values were computed using a 2-tailed t-test. (J-K’’’’) Larvae of the genotype CasExpress/G-trace; GAL80^ts^/+ were treated as in Fig. 1A. Wing discs were dissected at 20-22h after irradiation with 4000R, fixed and stained with an antibody against γ-H2Av. The discs were also stained for DNA and imaged for GFP. Arrowheads in J’’’’ indicate rare cells in which GFP and γ-H2Av signals overlap to appear white. The boxed area in J’’’ is magnified in K-K’’’’. L-L’’’ show a magnified area of another disc, to demonstrate reproducibility. A total of 52 discs were examined in 3 biological replicate experiments. Scale bar = 960 microns in A-F, 120 microns in J and 48 microns in K-L.

There are two possible interpretations of the data that cells with XPAC show reduced LOH. Non-lethal caspase activity may facilitate DNA repair to reduce genome instability or non-lethal caspase activity and chromosome breaks may act in a synthetic lethal combination to take the cell above the threshold for apoptosis, thereby eliminating such cells. To distinguish between these possibilities, we monitored DNA repair kinetics after irradiation and correlated them with non-lethal caspase activity. γ-H2Av signal (fly γ-H2Ax) is detected in all cells of the wing disc after exposure to 4000R of X-rays indicative of DNA double strand breaks induction (4h timepoint shown in S1 Figure) but returns to background levels 24h after IR as repair is completed (18, 24). Consistent with these data, at 20-22h after IR, only a subset of cells show γ-H2Av staining (Fig 6 J-L’’’’), which we interpret as cells that have yet to complete DNA repair. CasEx>GFP, which becomes detectable at similar times, show mutual exclusion with γ-H2Ax; cells that show both are rare (arrowheads in Fig 6J’’’’). We interpret these results to mean that XPAC cells complete DNA repair more efficiently than their GFP-counterparts. More efficient repair in XPAC cells, we propose, could explain the observed reduction in LOH in the adult wing.

## Discussion

We report here a study of cells with X-ray induced past active caspases (XPAC) during wing development in *Drosophila*. An optimized temperature shift protocol allowed us to minimize cells with spontaneous, developmental caspase activity and to focus specifically on XPAC cells. Such cells, we found, were able to proliferate and contribute to the adult organ and show reduced genome aberrations as measured by LOH, which results primarily from aneuploidy of whole or segments of chromosomes in *Drosophila* (29). In mouse embryo fibroblasts and immortalized human epithelial cells and fibroblasts (MCF10A and IMR90) exposed to ^56^F ion particle radiation, caspase activity, γ-H2Ax signal and DNA fragmentation persisted for weeks (10). Irradiated MCF10A cells with high caspase reporter activity formed colonies in soft agar compared to irradiated cells with low reporter activity. Inhibition of caspase 3 using shRNA or a dominant negative construct reduced the colony number, suggesting that caspase activity was causing transformation. While contrary to the findings in mammalian cells, our LOH results are consistent with the wide-spread non-lethal caspase activity during development in *Drosophila* (11, 12) and other metazoans. Clearly, caspases activity in these cases is not causing DNA fragmentation or it would have been detected in multiple studies that monitored DNA breaks (e.g., (18, 24)). We interpret our data to mean that non-lethal caspase activity protects the genome from X-ray-induced chromosome breaks.

Unlike with ^56^F radiation, X-ray-induced DSBs are cleared within 24h, even with much higher IR doses of 4000R (40 Gy) (18, 24). DNA repair may be more efficient in wing disc cells and that in this context, non-lethal caspase activity is beneficial. The question is, how? Caspase-driven cellular transformation in the above cited studies was proposed to occur because caspases work through endonuclease G to unravel the genome to allow access to oncogenic transcription factors such as KRAS/E1A used in these studies (10). Likewise, caspases could make the genome more accessible to DNA repair enzymes instead of transformation factors in the wing disc. In other words, consequences of non-lethal caspase activity may depend on the cellular context, specifically, which proteins are present and active to access the DNA. In cells predisposed to transformation such as immortalized cell lines, the outcome may be transformation. In cells that lack transformation factor but are robustly expressing DNA repair enzymes, improved repair may be the outcome.

A prior study used the CasEx>GFP reporter and found that Dronc and p35 affect developmental non-lethal caspase activity (14), similar to what we see for XPAC, and identified Akt and transcription factor dCIZ1 as new regulators of developmental CasEx>GFP induction. One striking difference, however, concerns the effect of oncogenic Ras. It preserves cells with developmental caspase activation; expression of Ras^V12^ in the posterior compartment of wing discs <u>increased</u> CasEx>GFP area by 2-fold. In contrast, Ras^V12^ <u>decreased</u> the XPAC area by 2-fold (Fig 5F), which we attribute to its ability to prevent caspase amplification like Wg, Dronc^DN^ and p35 (Fig. 3H). We conclude that retention of cells with past caspase activity shows common as well as different regulatory controls depending on whether cells are subject to developmental or IR-induced caspase activity.

## MATERIALS AND METHODS

### Drosophila stocks

*Drosophila* stocks are listed in S1 Table and include Caspase Tracker (12), UAS-hid/rpr (30), UAS-Wg (31) and UAS-Dronc^DN^ (32). The following stocks were generated using standard *Drosophila* techniques:

Caspase Tracker/CyO-GFP; GAL80^ts^/GAL80^ts^

Caspase Tracker/CyO-GFP; mwh^1^/TM6 Tb

G-trace/CyO-GFP; UAS-X/TM6 Tb where X was UAS-Wg, UAS-Axin, UAS-p35, UAS-Dronc^DN^, UAS-Ras^V12^, UAS-Notch^ICD^, UAS-mam^Δ^, or H99

G-trace/CyO-GFP; GAL80^ts^/GAL80^ts^

UAS-hid, UAS-rpr/UAS-hid, UAS-rpr or Y; G-trace/CyO-GFP

UAS-rpr/UAS-rpr or Y; G-trace/CyO-GFP

### Larvae culture, temperature shift and irradiation

After crossing virgins and males, the adults were fed on Nutri-Fly German Formula food (Genesee Scientific) for 3 days at 25°C to boost egg production, before transferring them to 18°. Embryos were collected and larvae were raised on Nutri-Fly Bloomington Formula food (Genesee Scientific). The cultures were monitored daily for ‘dimples’ on the food surface that denotes crowding and split as needed. Larvae in food were placed in petri dishes and irradiated at room temperature (rt) in a Faxitron Cabinet X-ray System Model RX-650 (Lincolnshire, IL) at 115 kV and 5.33 rad/sec. Irradiated larvae in food were placed in new vials in temperature bath filled with Lab Armor beads (ThermoFisher), which we found gave more consistent and reproducible data than water baths. After the temperature shift to inactivate GAL80, larvae were returned to 18°C until dissection.

### Tissue preparation, staining and imaging

For Acridine Orange staining, Larvae were dissected in PBS, and imaginal discs were incubated for 5 min in PBS+0.5 mM AO (Sigma) at rt, washed twice with PBS, mounted in PBS, and imaged immediately. For γ-H2Av staining, wing discs were dissected in PBS, fixed in 2% paraformaldehyde (PFA) in PBS for 45 min, washed twice with PBS and permeabilized in PBS+0.5% Triton X-100 for 12.5 min. For other antibodies, wing discs were dissected in PBS, fixed in 4% PFA in PBS for 30 minutes, washed thrice in PBS, permeabilized in PBS+0.5% Triton X-100 for 10 min, and rinsed in PBTx (0.1% Triton X-100). The discs were blocked in 5% Normal Goal Serum in PBTx for at least 1 h and incubated overnight at 4°C in primary antibody in block. Primary antibodies were to cleaved Dcp1 (1:100, rabbit polyclonal, Cell Signaling Cat# 9578), Phospho-Histone H3 (1:1000, rabbit monoclonal, Upstate Biotech), Wg (1:100, mouse monoclonal, DSHB#4D4) and γ-H2Av (1:1000, mouse monoclonal, DSHB#UNC93-5.2.1). The discs were rinsed thrice in PBTx and blocked for at least 30 min before incubation in secondary antibody in block for 2 h at room temperature. Secondary antibodies were used at 1:400-500 (Jackson). The discs were washed in PBTx and stained with 10 ug/ml Hoechst33258 in PBTx for 2 min, washed 3 times, and mounted on glass slides in Fluoromount G (SouthernBiotech).

For adult wings, newly eclosed adults with unfurl wings were collected and kept at rt. The wings were collected within 30 min of unfurling, submerged in PBTx, mounted in Fluoromount G (SouthernBiotech), and imaged immediately. Any delay meant the loss of fluorescence as wing epithelial cells are eliminated in a developmentally regulated wing maturation step [28, 29].

Wing discs and wings were imaged on a Leica DMR compound microscope using a Q-Imaging R6 CCD camera and Ocular or Micro-Manager software. Images were processed and analyzed using ImageJ software.

### Statistical Analysis

For sample size justifications, we used a simplified resource equation from (33); E = Total number of animals − Total number of groups, where E value of 10-20 is considered adequate. When we compare two groups (e.g., -/+IR), n= 6 per group or E = 11 would be adequate. All samples exceed this criterion. 2-tailed Student t-test with equal variance was used.

## Conflict of Interest

Authors declare no competing financial interests in relation to the work described.

## Acknowledgements (include funding sources)

*Drosophila* stocks from the Bloomington *Drosophila* Stock Center (NIH P40 OD018537) were used in this study. The authors thank Hardwick and Tang labs for Caspase Tracker stocks, Montell lab for CasExpress stocks, and Kristin White for the critical reading of the manuscript. This work was funded by an NIH grant and a diversity supplement to TTS (R35 GM130374) and an NIH fellowship to SCP (F31 GM149184). Monoclonal antibodies developed by Joshua Sanes, Stephen Cohen, and R. Scott Hawley were obtained from the Developmental Studies Hybridoma Bank, created by the NICHD of the NIH and maintained at The University of Iowa, Department of Biology, Iowa City, IA 52242.

## Figure Legends

**S1 Figure.**
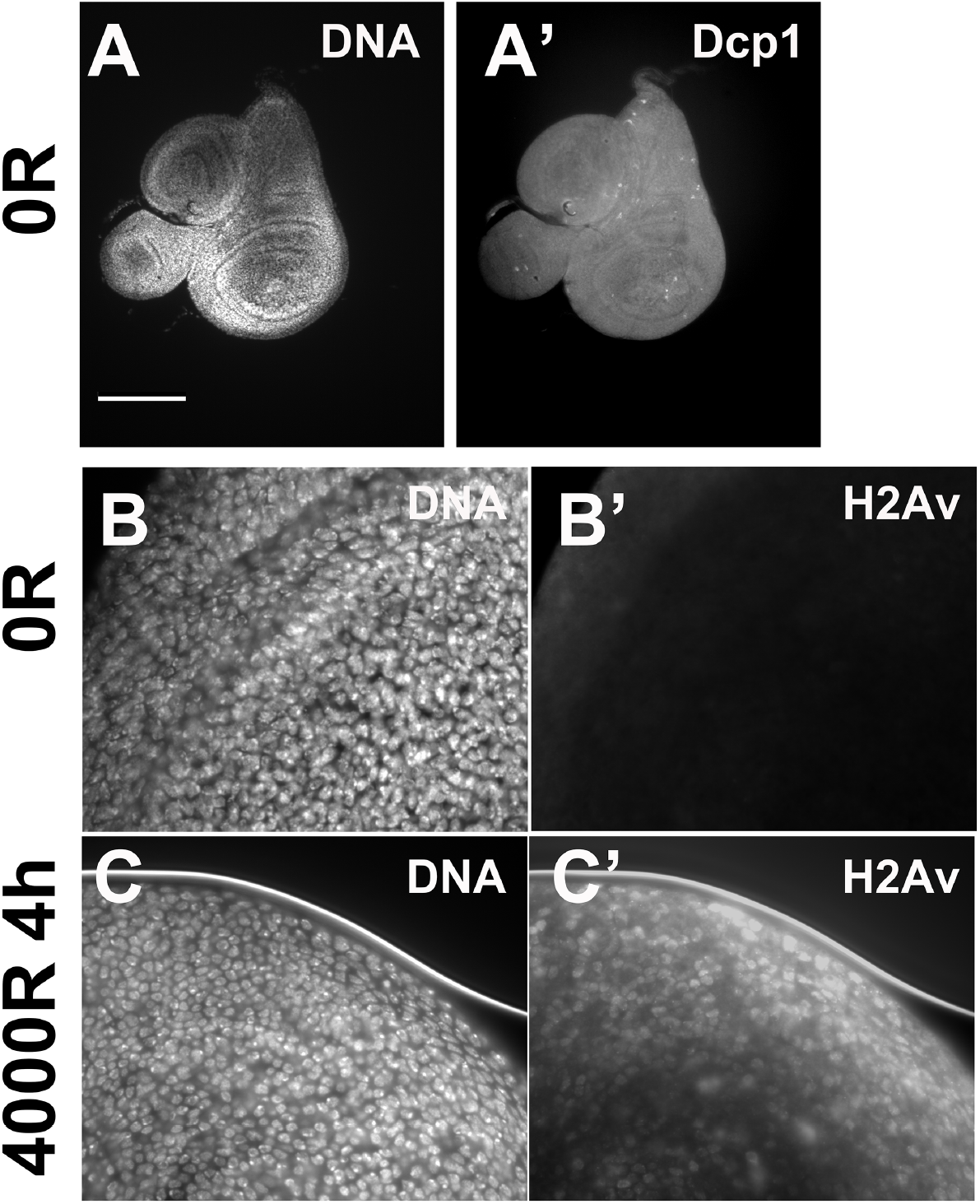
Caspase cleavage without IR and DNA double strand breaks with and without IR. Larvae of the genotype CasExpress/G-trace; GAL80ts/+ were treated as in Fig. 1A where T=26°C. Wing discs were dissected at 4h after exposure to 0 or 4000R of X-rays, fixed, and stained for DNA and with antibodies to cleaved Dcp1 (A-A’) or γ-H2Av (B-C’). The scale bar = 120 microns in A and 24 microns in B-C.

**S1 Table.**
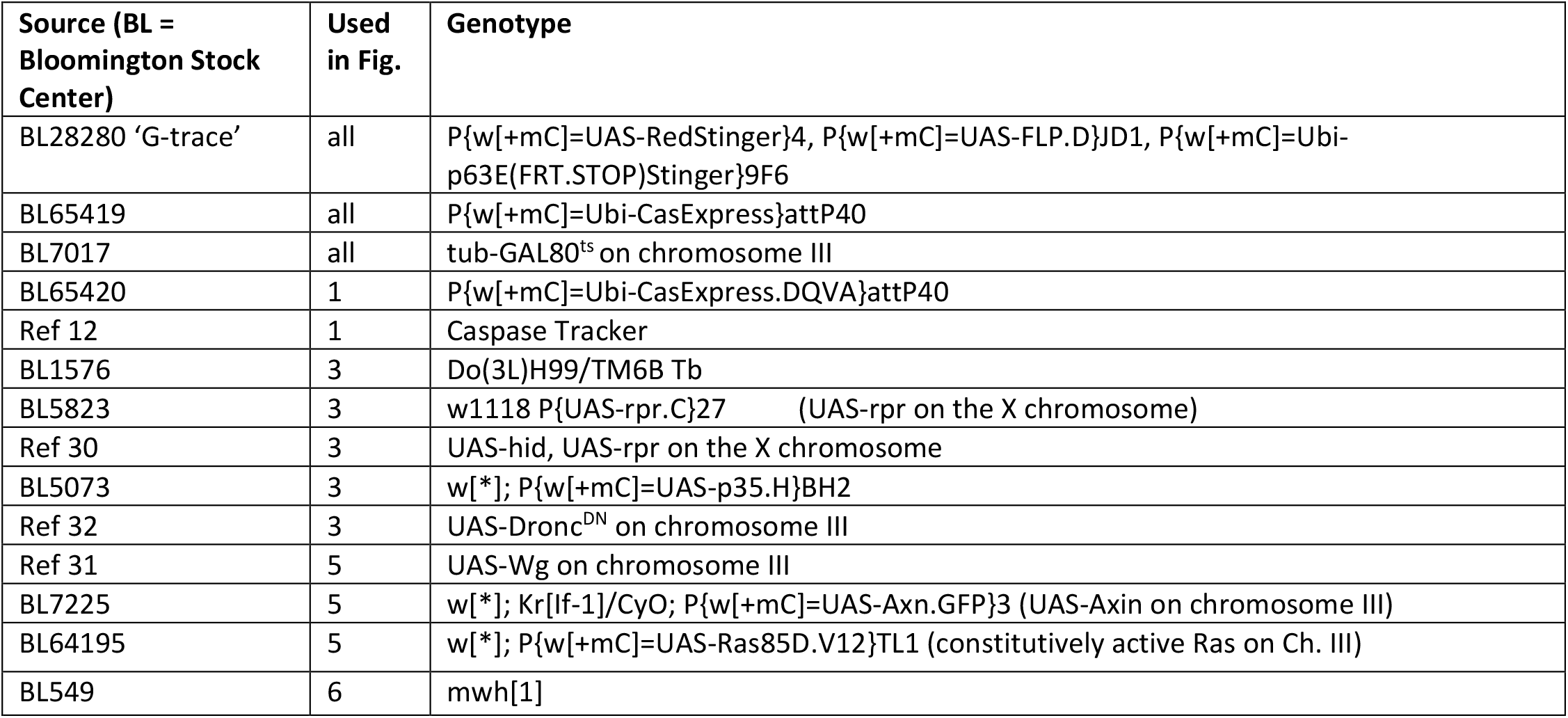
Fly stocks used.

## Notes

### Competing Interest Statement

The authors have declared no competing interest.

### Summary of Updates

We have shortened the text to fit the journal format and have added new data.

## References

1. Chakraborty S, Mir KB, Seligson ND, Nayak D, Kumar R, Goswami A. Integration of EMT and cellular survival instincts in reprogramming of programmed cell death to anastasis. Cancer Metastasis Rev. 2020;39(2):553–66.

2. Gudipaty SA, Conner CM, Rosenblatt J, Montell DJ. Unconventional Ways to Live and Die: Cell Death and Survival in Development, Homeostasis, and Disease. Annu Rev Cell Dev Biol. 2018;34:311–32.

3. Su TT. Non-apoptotic roles of apoptotic proteases: new tricks for an old dog. Open Biol. 2020;10(8):200130.

4. Tang HM, Tang HL. Anastasis: recovery from the brink of cell death. R Soc Open Sci. 2018;5(9):180442.

5. McArthur K, Kile BT. Apoptotic Caspases: Multiple or Mistaken Identities? Trends Cell Biol. 2018;28(6):475–93.

6. Nakajima YI, Kuranaga E. Caspase-dependent non-apoptotic processes in development. Cell Death Differ. 2017;24(8):1422–30.

7. Tang HL, Tang HM, Mak KH, Hu S, Wang SS, Wong KM, et al. Cell survival, DNA damage, and oncogenic transformation after a transient and reversible apoptotic response. Mol Biol Cell. 2012;23(12):2240–52.

8. Tang HM, Talbot CC, Jr., Fung MC, Tang HL. Molecular signature of anastasis for reversal of apoptosis. F1000Res. 2017;6:43.

9. Ichim G, Lopez J, Ahmed SU, Muthalagu N, Giampazolias E, Delgado ME, et al. Limited mitochondrial permeabilization causes DNA damage and genomic instability in the absence of cell death. Mol Cell. 2015;57(5):860–72.

10. Liu X, He Y, Li F, Huang Q, Kato TA, Hall RP, et al. Caspase-3 promotes genetic instability and carcinogenesis. Mol Cell. 2015;58(2):284–96.

11. Ding AX, Sun G, Argaw YG, Wong JO, Easwaran S, Montell DJ. CasExpress reveals widespread and diverse patterns of cell survival of caspase-3 activation during development in vivo. Elife. 2016;5.

12. Tang HL, Tang HM, Fung MC, Hardwick JM. In vivo CaspaseTracker biosensor system for detecting anastasis and non-apoptotic caspase activity. Sci Rep. 2015;5:9015.

13. Evans CJ, Olson JM, Ngo KT, Kim E, Lee NE, Kuoy E, et al. G-TRACE: rapid Gal4-based cell lineage analysis in Drosophila. Nat Methods. 2009;6(8):603–5.

14. Sun G, Ding XA, Argaw Y, Guo X, Montell DJ. Akt1 and dCIZ1 promote cell survival from apoptotic caspase activation during regeneration and oncogenic overgrowth. Nat Commun. 2020;11(1):5726.

15. Verghese S, Su TT. Ionizing radiation induces stem cell-like properties in a caspase-dependent manner in Drosophila. PLoS Genet. 2018;14(11):e1007659.

16. Wichmann A, Jaklevic B, Su TT. Ionizing radiation induces caspase-dependent but Chk2- and p53-independent cell death in Drosophila melanogaster. Proc Natl Acad Sci U S A. 2006;103(26):9952–7.

17. Abrams JM, White K, Fessler LI, Steller H. Programmed cell death during Drosophila embryogenesis. Development. 1993;117(1):29–43.

18. Wells BS, Johnston LA. Maintenance of imaginal disc plasticity and regenerative potential in Drosophila by p53. Dev Biol. 2012;361(2):263–76.

19. Neufeld TP, de la Cruz AF, Johnston LA, Edgar BA. Coordination of growth and cell division in the Drosophila wing. Cell. 1998;93(7):1183–93.

20. Tozluoglu M, Duda M, Kirkland NJ, Barrientos R, Burden JJ, Munoz JJ, et al. Planar Differential Growth Rates Initiate Precise Fold Positions in Complex Epithelia. Dev Cell. 2019;51(3):299–312 e4.

21. McComb S, Chan PK, Guinot A, Hartmannsdottir H, Jenni S, Dobay MP, et al. Efficient apoptosis requires feedback amplification of upstream apoptotic signals by effector caspase-3 or -7. Sci Adv. 2019;5(7):eaau9433.

22. Fujita E, Egashira J, Urase K, Kuida K, Momoi T. Caspase-9 processing by caspase-3 via a feedback amplification loop in vivo. Cell Death Differ. 2001;8(4):335–44.

23. Florentin A, Arama E. Caspase levels and execution efficiencies determine the apoptotic potential of the cell. J Cell Biol. 2012;196(4):513–27.

24. Verghese S, Su TT. Drosophila Wnt and STAT Define Apoptosis-Resistant Epithelial Cells for Tissue Regeneration after Irradiation. PLoS Biol. 2016;14(9):e1002536.

25. Johnston LA, Edgar BA. Wingless and Notch regulate cell-cycle arrest in the developing Drosophila wing. Nature. 1998;394(6688):82–4.

26. Bergmann A, Agapite J, McCall K, Steller H. The Drosophila gene hid is a direct molecular target of Ras-dependent survival signaling. Cell. 1998;95(3):331–41.

27. Lu Q, Schafer DA, Adler PN. The Drosophila planar polarity gene multiple wing hairs directly regulates the actin cytoskeleton. Development. 2015;142(14):2478–86.

28. Brown J, Bush I, Bozon J, Su TT. Cells with loss-of-heterozygosity after exposure to ionizing radiation in Drosophila are culled by p53-dependent and p53-independent mechanisms. PLoS Genet. 2020;16(10):e1009056.

29. Baker BS, Carpenter AT, Ripoll P. The Utilization during Mitotic Cell Division of Loci Controlling Meiotic Recombination and Disjunction in DROSOPHILA MELANOGASTER. Genetics. 1978;90(3):531–78.

30. Zhou L, Schnitzler A, Agapite J, Schwartz LM, Steller H, Nambu JR. Cooperative functions of the reaper and head involution defective genes in the programmed cell death of Drosophila central nervous system midline cells. Proc Natl Acad Sci U S A. 1997;94(10):5131–6.

31. Lawrence PA, Sanson B, Vincent JP. Compartments, wingless and engrailed: patterning the ventral epidermis of Drosophila embryos. Development. 1996;122(12):4095–103.

32. Verghese S, Bedi S, Kango-Singh M. Hippo signalling controls Dronc activity to regulate organ size in Drosophila. Cell Death Differ. 2012;19(10):1664–76.

33. Charan J, Kantharia ND. How to calculate sample size in animal studies? J Pharmacol Pharmacother. 2013;4(4):303–6.

